# Endogenous DAF-16 Spatiotemporal Activity Quantitatively Predicts Lifespan Extension Induced by Dietary Restriction

**DOI:** 10.1101/2021.12.20.473576

**Authors:** Javier Huayta, Adriana San-Miguel

## Abstract

In many organisms, dietary restriction (DR) leads to lifespan extension through the activation of cell protection and pro-longevity gene expression programs. In the nematode *C. elegans*, the DAF-16 transcription factor is a key aging regulator that governs the Insulin/IGF-1 signaling pathway and undergoes translocation from the cytoplasm to the nucleus of cells when animals are exposed to food limitation. In this work, we assess the endogenous activity of DAF-16 under various DR regimes by coupling CRISPR/Cas9-enabled fluorescent tagging of DAF-16 with quantitative image analysis and machine learning. Our results indicate that lifelong DAF-16 endogenous activity is a robust predictor of mean lifespan in *C. elegans*, and it accounts for 78% of the lifespan variability induced by DR. We found that this lifespan-extending mechanism occurs mainly in the intestine and neurons, and that DR drives DAF-16 activity in unexpected locations such as the germline and intestinal nucleoli.

## Introduction

*C. elegans* exposed to plentiful food and a stress-free environment can sustain growth and reproduction. Under harsh environmental conditions such as food depravation, oxidative and heat stress, and others, a number of genetic pathways activate to promote cell protection and survival in *C. elegans*. These pathways are governed by transcription factors such as DAF-16, PHA-4, and SKN-1 (Kenyon, 2010). The best known of these environmental factors is dietary restriction (DR), which extends lifespan in many species (Masoro, Yu and Bertrand, 1982; Weindruch, 1996; Masoro, 2005; Mattison *et al*., 2017). In *C. elegans*, the DAF-16 transcription factor regulates the Insulin/IGF-1 signaling pathway. Under various forms of stress, DAF-16 migrates from the cytoplasm to the nucleus inducing lifespan extension (Lin *et al*., 2001) and attenuating the effects of aging in *C. elegans* (Hsu, Murphy and Kenyon, 2003). Nevertheless, it is still unclear how important is the magnitude of lifelong DR exposure and the responses of longevity-regulating genes in quantitatively determining lifespan in *C. elegans*. Previous work has shown that these genetic pathways can be manipulated to modulate gene expression and extend lifespan (Sagi and Kim, 2012; Win *et al*., 2013; Stroustrup *et al*., 2016). Furthermore, systems approaches have been used to identify regulatory networks of the DR response, which can be targeted to manipulate the effects of DR (Hou *et al*., 2016). However, whether the spatiotemporal activity of these transcription factors is sufficient to predict the lifespan extension conveyed by their stress-induced activities is still unclear. Elucidating the contributions of these components to *C. elegans* survival, would permit interventions that target lifespan augmentation, and shed light on the relative importance of different pathways towards longevity. With these considerations, we analyze lifespan in *C. elegans* as the outcome of cumulative molecular activity of DAF-16 prompted by exposure to DR.

Various approaches have been successfully used to probe gene expression in *C. elegans* cells, tissues, and at different developmental stages, such as DNA microarrays, real-time PCR, RNAseq, and serial analysis of gene expression (SAGE) (McElwee, Bubb and Thomas, 2003; Portman, 2006; Wang *et al*., 2009; Guthmueller, Yoder and Holgado, 2011; Possik and Pause, 2015). These methods are destructive in nature, as they require extraction of DNA and RNA. Therefore, these techniques are not suitable to obtain spatial information of the translocation of cytoplasmic DAF-16 protein to nuclei. Previous studies have also used strains with fluorescently tagged DAF-16, which carry transgenic arrays that typically have hundreds of copies of the gene (Stinchcomb *et al*., 1985) and label only specific isoforms (Libina, Berman and Kenyon, 2003; Kwon *et al*., 2010; Kumsta and Hansen, 2012). Here, we measure endogenous activity of all DAF-16 isoforms by tagging the *daf-16* locus at the 3’ end using CRISPR/Cas9 (Dickinson *et al*., 2015). Coupled with spatiotemporal quantification of nuclear fluorescence through quantitative image analysis and machine learning, this approach enables *in vivo* analysis of all DAF-16 activity in *C. elegans* at endogenous levels, in a tissue-specific manner. This approach reveals that DAF-16 spatiotemporal activity (measured as lifelong total nuclear intensity) is a strong predictor of lifespan in *C. elegans* populations exposed to DR in liquid culture. Furthermore, we show that the main contributors to this DAF-16 activity are neurons and intestinal cells, indicating that the observed lifespan extension originates mainly from the translocation and activity of DAF-16 in these types of cells. Finally, we demonstrate that DAF-16 activity is observed in unexpected locations such as germ cells and intestinal nucleoli, where DAF-16 could be undergoing yet-to-be described interactions affecting the aging process.

## Methods

### Strains, media, and culture

*C. elegans* was maintained on standard Nematode Growth Medium (NGM) plates seeded with OP50 *E. coli* bacteria and kept at 20 ºC until they started to lay eggs. Animals were then bleached using standard protocols to obtain age-synchronized populations (Porta-de-la-Riva *et al*., 2012). This process was repeated 3 times in total to avoid transgenerational epigenetic effects related to longevity (Greer *et al*., 2011). L4 animals were then transferred to a cell culture flask containing 4ml of SB media with 10^10^ cells/ml of OP50 bacteria and 100 μM 5-fluorodeoxyuridine (FUdR) and kept at 20 ºC for 1 day (Stiernagle, 2006; Gruber *et al*., 2009). Cultures were then used for dietary restriction and lifespan experiments. Strains used in this work were ASM10 *daf-16* (del2 [*daf-16*::GFP-C1^3xFlag]) I and N2 (*C. elegans* wild isolate). OP50 *E. coli* was grown in LB media following standard procedures (Amrit *et al*., 2014). Bacteria were washed thrice with SB media, pelletized, and suspended in S-Medium at a concentration of 100 mg/mL, corresponding to a concentration of 2 × 10^10^ cells/ml.

### Generation of transgenic line

We used a CRISPR/Cas9 approach developed by Dickinson *et al*. to insert a GFP-encoding sequence at the 3’ end of *daf-16* (Dickinson *et al*., 2015). We chose the Cas9 target site by identifying all possible single guide RNAs (sgRNA) in a 200bp region centered in the stop codon of the target gene. This region was selected to permit introduction of the fluorescent tag at the C-terminus, tagging in this manner all possible isoforms of the gene. Criteria such as specificity, activity, and distance to the stop codon were considered in selecting the sgRNA using the GuideScan design tool (Perez *et al*., 2017). This 20bp sequence was introduced into a Cas9-encoding construct (pDD162, Addgene #47549) using a NEB Q5 Site-Directed Mutagenesis Kit to generate plasmid pJHR1. 500-700bp long homology arms at each side of the *daf-16* stop codon were then generated by PCR amplification of genomic DNA from N2 animals. These fragments were isolated, purified, and introduced in a construct containing a selection cassette (pDD282, Addgene #66823) using NEBuilder HiFi DNA Assembly Mix to generate plasmid pJHR2 (**Table 1, Figure S1, S2**). A mix containing 15ng/μL pJHR1, 50ng/μL pJHR2, and 2.5ng/μL pCFJ90 (mCherry co-injection marker) was injected in the gonads of 100 N2 young-adult animals following standard microinjection procedures (Evans, 2006). These animals were transferred to NGM plates and left to produce progeny at 25 °C. Hygromycin was added to the plates to kill untransformed F1 progeny. After 7 days, animals showing the Rol phenotype and lacking red fluorescent extrachromosomal array markers were transferred to new NGM plates without hygromycin. Individual putative knock-in animals were transferred to new plates for homozygous selection. Finally, L1 larvae from homozygous plates (those that contained only animals with the Rol phenotype) were heat-shocked at 34 °C for 4 hours to induce Cre expression, and excision of the selection cassette. Wild-type adult animals lacking the Rol phenotype were picked and maintained as the strain containing the fluorescent-tagged gene. PCR and Sanger sequencing were performed to confirm correct introduction of the tag.

### Dietary restriction and fluorescence imaging

Approximately 200 worms in day 1 of adulthood kept in liquid culture were washed 3 times with SB media and moved to a flask containing 4ml of SB media with a reduced food concentration to induce DR (10^9^, 10^8^ or 0 OP50 cells/ml) for a fixed amount of time (6, 12, or 24 hours). After the DR period concluded, additional OP50 was added to raise the concentration to an *ad libitum* level (10^10^ OP50 cells/ml). FUdR concentration was raised to 100 μM total concentration to prevent offspring production (Mitchell *et al*., 1979). Worms were then kept at 20 ºC for 3 days at *ad libitum* food concentration. DR exposure was repeated on days 4, 7, and 10 of adulthood. At the start and end of each DR exposure, approximately 20-30 worms were immobilized using 10 μL of 2 mM tetramisole on dried 2% agarose pads of approximately 1 cm diameter. Confocal fluorescence microscopy was performed with a Leica DMi8 microscope paired with an 89 North LDI Laser Diode Illuminator coupled with a CrestOptics X-Light V2 confocal imager and an Orca-Flash 4.0 digital CMOS camera. The exposure time and laser intensity were kept constant throughout all experiments at 100 ms and 100%, respectively. Images containing 30 slices (Z-stacks with 3 μm spacing) were acquired with a MATLAB GUI (Graphical User Interface). Images were acquired in the green and red channels sequentially. This enables later subtraction of autofluorescence present in both channels with a custom MATLAB script.

### Lifespan experiments

Approximately 200 worms per flask were kept in liquid culture in the same conditions described in the previous section. Animal mobility was analyzed in a stereoscope through visual inspection. Nematodes were scored as alive (active and swimming), dead (rigid and not moving), and censored (lost to manipulation). This was performed every third day, starting on day 1 of adulthood. Lifespan curves were constructed using Online Application for Survival Analysis 2 (Han *et al*., 2016) to obtain the mean lifespan for each population.

### Quantitative image processing

Images were analyzed using a MATLAB script, customized to segment cell nuclei in each slice of the z-stack. The algorithm’s most important steps are: 1) Subtraction of red channel image from green channel image to eliminate autofluorescence present in both images; 2) Generation of a binary mask from the subtracted image; 3) Identification of nuclei by retaining segmented objects that: a) filled up at least 50% of their bounding box, b) had eccentricity values under 0.85, and c) had solidity values above 0.75. The threshold values were determined based on visual assessment of the characteristics of nuclei. Object characteristics were extracted using “regionprops”; 4) Use of the “imopen” function to smooth edges. 5) Deletion of objects smaller than neurons and larger than intestinal cells using the “bwareaopen” function. These operations enable for the selection of nuclei (ellipse or circle-like objects) while eliminating other features such as autofluorescence, animal edges, embryos, etc. in a stepwise manner (**Figure 1A**). To avoid over-estimation, nuclei present in multiple slices were identified by comparing their centroids. If multiple centroids with similar values in the ± 4 pixels range were identified in multiple slices, only the object with the largest area was retained for properties extraction. The MATLAB regionprops function was then used to quantify the total intensity in the segmented nuclei and the data extracted was stored in a MATLAB nested structure. Total intensity was calculated as the sum of the intensity of the pixels from all cell nuclei extracted per worm.

**Figure 1:**
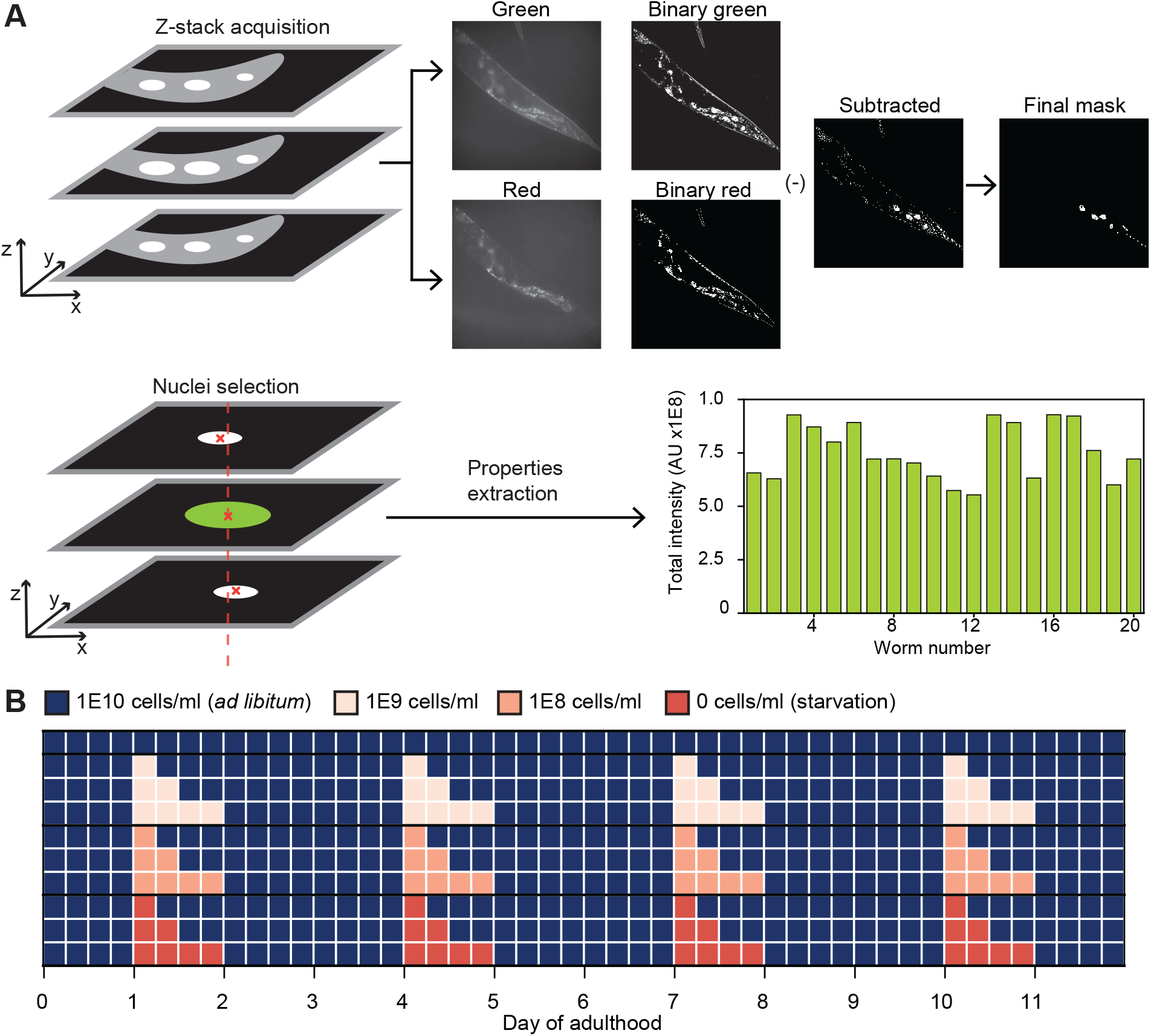
Quantitative analysis of endogenous DAF-16 under dietary restriction regimes. A) Four Z-stacks of 30 slices are taken of each animal. Each slice in the green channel has a corresponding red channel slice. Both are converted to binary images and subtracted from each other to eliminate autofluorescence. The resulting image is treated with morphological functions and objects are filtered based on extent, solidity, and eccentricity. An image containing only cell nuclei is obtained, and largest nuclei are retained if they appear in several slices. Pixel-based intensity is then aggregated to obtain total DAF-16 intensity in all of the animal’s identified nuclei. B) Diagram of DR regimes tested in this study. Animals were exposed to three levels of DR: 1E9, 1E8, or 0 OP50 cells/ml; for three exposure times: 6, 12, or 24 hours. After each exposure, animals were put back in an *ad libitum* concentration (1E10 OP50 cells/ml). Exposure to DR was performed at days 1, 4, 7, and 10 of adulthood.

### Cell type classification and nucleolus analysis

To classify nuclei belonging to specific tissues, we used a Fine Tree multi-class classification algorithm. A ground truth set of 600 cell nuclei and their corresponding class (intestine, neuron, hypodermis, or muscle) was manually generated. The Fine Tree algorithm was trained using MATLAB Classification Learner, by using the following object properties as features: area, eccentricity, and equivalent diameter. The generated Fine Tree algorithm resulted in a 96.1% accuracy when predicting cell types with a validation set.

For nucleolar analysis, DAF-16-filled nucleoli in intestinal cells were counted manually using FIJI. A nucleolus was considered “empty” when the nucleus’ center was empty. A nucleolus was considered “filled” when the nucleus center had fluorescence due to GFP-tagged DAF-16.

### Data analysis and statistics

Linear regressions were performed in Origin 2020b using the FitLinear function. Polynomial fits were performed in Origin 2020b using the FitPolynomial function. Significance tests such as t-Test and ANOVA were performed using the Data Analysis Add-on in MS Excel 2016. Lifespan analysis was performed with the Online Application for Survival Analysis 2 (Han *et al*., 2016) by using number of dead and censored animals as input; obtaining mean lifespan with a Kaplan-Meier estimator.

## Results

### DR regimes modulate endogenous DAF-16 activity

To study the quantitative link between endogenous DAF-16 activity, and lifespan, we first aimed to analyze the response of endogenous DAF-16 to multiple DR regimes by varying food concentration and exposure time. Using CRISPR/Cas9, we generated a strain with the endogenous *daf-16* locus labeled with GFP at the 3’ end, and measured the response of DAF-16 using a custom image processing algorithm that quantifies nuclear intensity throughout the entire animal. Accurate analysis of DAF-16::GFP levels is challenging, as single copy reporters lead to extremely dim images. Since GFP intensities from DAF-16 were lower or similar than those from autofluorescent lipid droplets, we performed image acquisition in the green and red channels (**Figure 1A**), allowing subtraction of autofluorescence from the GFP signal. Analysis was performed on every slice of a z-stack covering the entire animal, and four separate z-stacks per channel were acquired per animal to cover the entire worm length. Information from the four z-stacks were aggregated for each worm.

Animals were exposed to DR conditions at day 1 of adulthood and every third day thereafter (**Figure 1B**). All DR conditions induced an increase in the total nuclear intensity of DAF-16 compared to well-fed animals used as control (**Figure 2A, B**). The strongest DAF-16 response was observed for intermediate DR exposure times, peaking at 12 hours. The longest exposure (24 hours) showed a higher response than the control, but not as pronounced as 6 and 12 hours. Longer DR exposures could result in malnutrition, and lead to reduced DAF-16 responsiveness. Alternatively, DAF-16 could be translocating from nuclei back to the cytoplasm, a phenomenon that has been previously observed during long-term starvation (Weinkove *et al*., 2006). We observed a similar pattern when varying food concentration. While all DR concentrations induce a response in DAF-16, the maximum DAF-16 response is observed for a mid-level concentration (10^8^ OP50 cells/ml) (**Figure 2B**). This non-monotonic response to food concentration has been previously observed (Henderson and Johnson, 2001; Greer *et al*., 2007), and our work suggests this is also true when varying DR exposure time. While prior work has shown that DAF-16 activity does not play a role in starvation-induced longevity when animals are cultured in solid media (Lee *et al*., 2006), our results suggest that starvation does induce DAF-16 activity in liquid media culture. This suggests that DAF-16 could be induced under starvation when additional conditions are met which differ from solid media, such as an increased energy consumption from swimming (Ghosh and Emmons, 2008; Laranjeiro *et al*., 2017).

**Figure 2:**
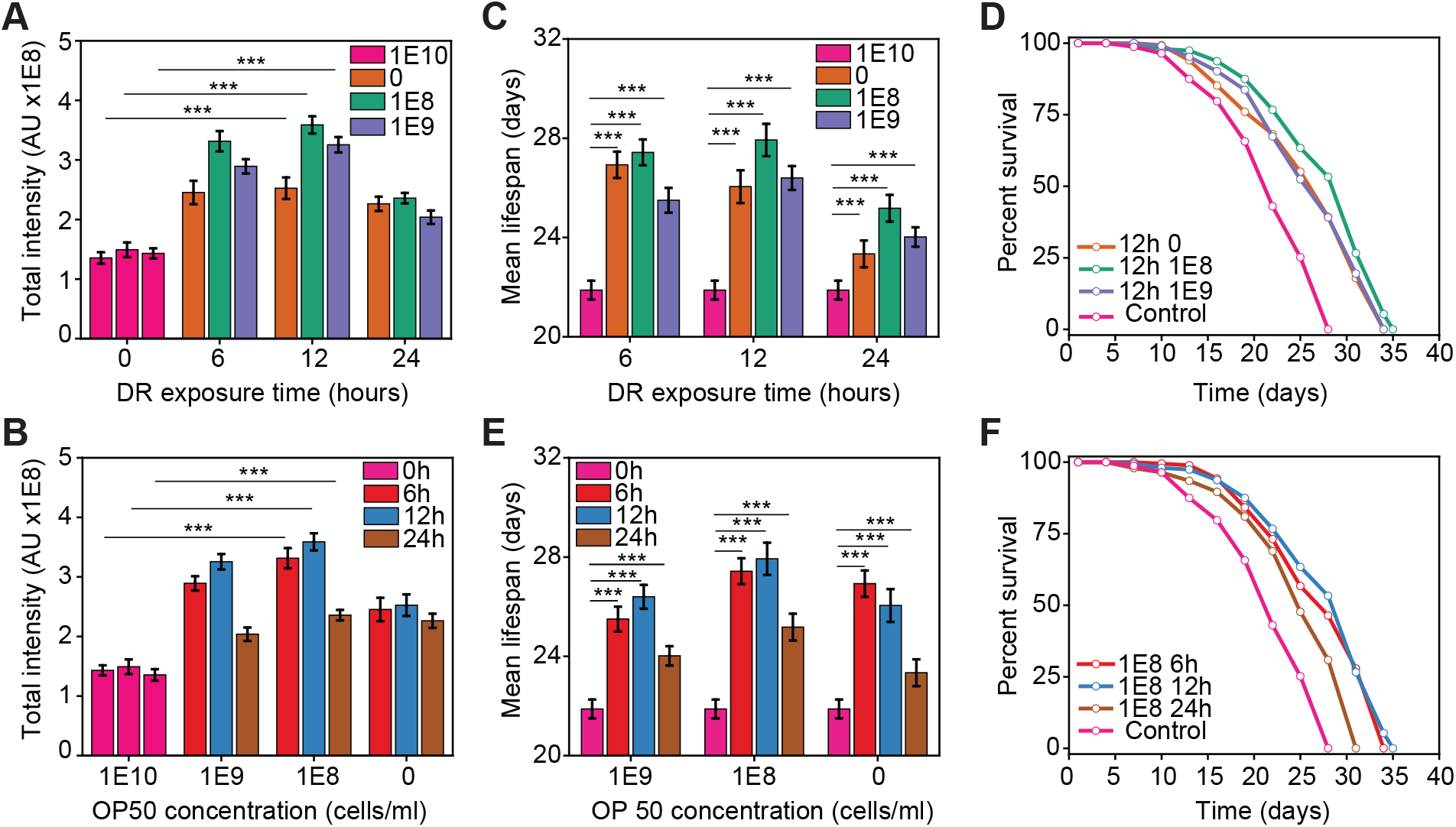
DR regimes modulate endogenous DAF-16 activity. A) DAF-16 total intensity as a function of exposure time to DR for various food concentrations at day 1 of adulthood. B) DAF-16 total intensity as a function of food concentration for various exposure times at day 1 of adulthood. C) Mean lifespan as a function of exposure time at various food concentrations. D) Lifespan curves for populations under various food concentrations with 12 hours of exposure to DR. E) Mean lifespan as a function of food concentration at various exposure times. F) Lifespan curves for populations under various exposure times to DR with a 10^8^ OP50 cells/ml food concentration. p < 0.001 (***). Error bars are SEM. All p-values were calculated using Tukey HSD for all pairwise comparisons after one-way ANOVA (unequal variances) comparison. Control population (pink) is *ad libitum* (1E10 cells/ml, 0 h exposure time).

To determine how important the magnitude of DAF-16 activity is in modulating lifespan, we measured mean lifespan under the different DR regimes mentioned previously. All DR regimes lead to an extension in mean lifespan when compared to the control (**Figure 2C-F, Figure S3**) as expected based on prior work (Kaeberlein *et al*., 2006; Greer and Brunet, 2009). In line with the results of DAF-16 activity, the largest lifespan extension was driven by the mid-level concentration of 10^8^ OP50 cells/ml (**Figure 2C, D, Figure S3**), and for 6 and 12 hours regimes (**Figure 2E, F, Figure S3**). The 24 hours regime also induced lifespan extension, but not as significantly as 6 or 12 hours (**Figure 2F**). It has been previously observed that long-exposure to starvation increases the likelihood of matricidal hatching, which leads to the animals’ death (Chen and Caswell-Chen, 2003). While we avoid this potential problem by exposing animals to FUdR, long-exposure to starvation could be inducing deleterious effects that are not fully abrogated by the beneficial pathways activated by DR in *C. elegans*.

### Cumulative lifelong DAF-16 nuclear activity determines lifespan under DR

We assessed the role of DAF-16 cumulative activity in lifespan extension by adding the total intensity of DAF-16 for all identified nuclei per animal at days 1, 4, 7, and 10 of adulthood, and comparing it to the corresponding mean lifespan (**Figure 3A**). We found that the cumulative activity of nuclear DAF-16 predicts mean lifespan with an R^2^ of 0.78 for the DR regimes explored in this study, using a linear regression. This suggests that lifespan extension due to DR is mostly modulated by cumulative DAF-16 lifelong spatiotemporal activity. The remainder of lifespan variability could come from DAF-16 activity at days we did not evaluate or, more likely, from the activity of other signaling pathways that also mediate longevity under DR, such as the target-of-rapamycin (TOR) pathway regulated by PHA-4 (Panowski *et al*., 2007; Hansen *et al*., 2008; Sheaffer, Updike and Mango, 2008) and the oxidative stress response regulated by SKN-1 (An and Blackwell, 2003; Bishop and Guarente, 2007).

**Figure 3:**
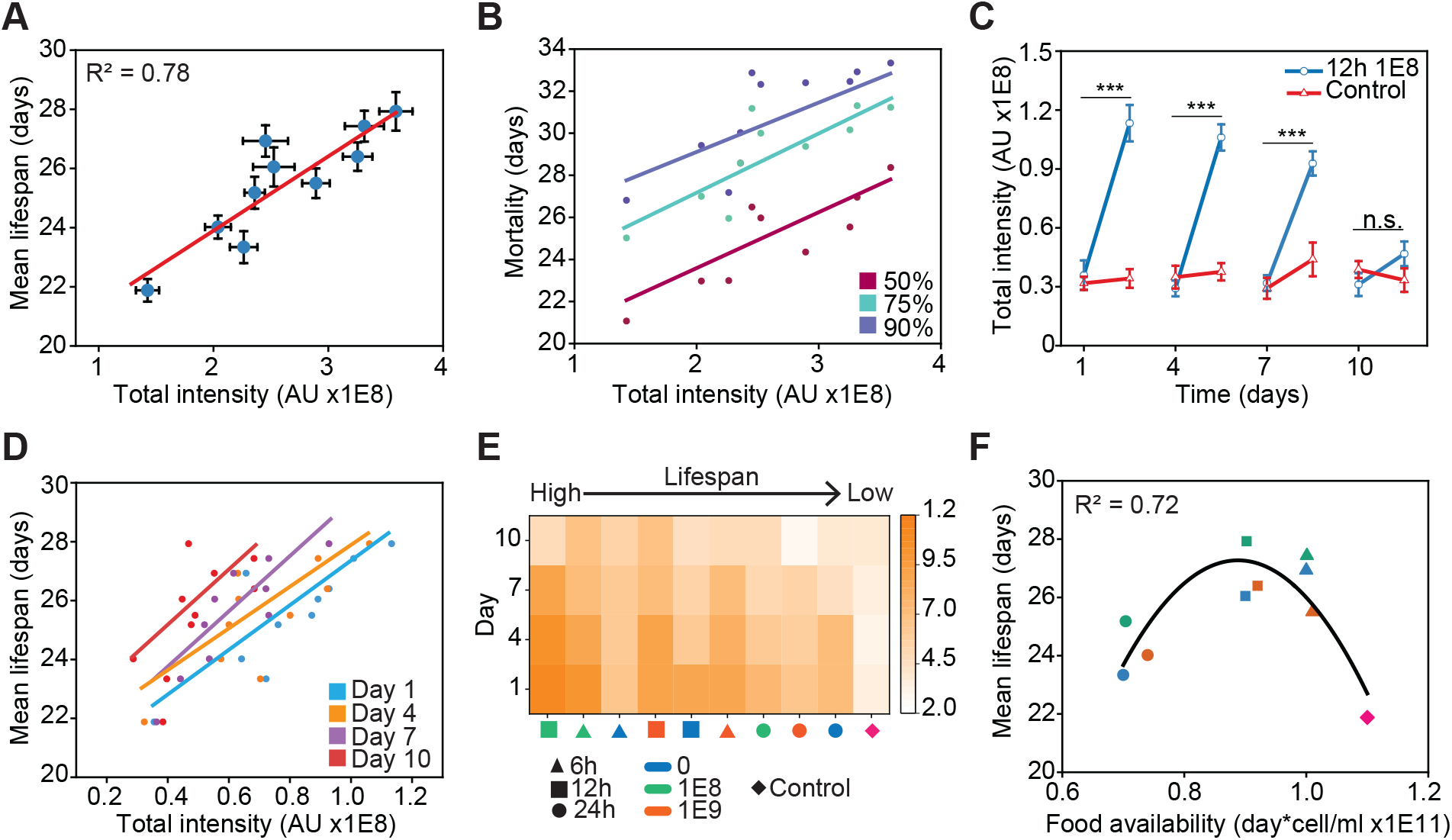
Cumulative lifelong DAF-16 nuclear activity determines lifespan. A) Mean lifespan of *C. elegans* as a function of lifelong DAF-16 total intensity under various DR regimes. B) Mortality of *C. elegans* populations as a function of DAF-16 lifelong total intensity. C) DAF-16 total intensity at a 10^8^ OP50 cells/ml food concentration with 12 hours of exposure time compared to an *ad libitum* control at 1, 4, 7, and 10 days of adulthood. D) Mean lifespan as a function of DAF-16 total intensity in individual days. E) Heat map of DAF-16 total nuclear intensity per day under different DR regimes, in order of higher to lower lifespan (left to right). F) Mean lifespan as a function of total food available during the first 11 days of animal adulthood. 6 hours (triangles) and 12 hours (squares) regimes are at the peak of the curve while 24 hours (circles) regimes and the control (diamond) are at the lower ends of the curve. Error bars are SEM. p > 0.05 (n.s.), p < 0.001 (***). All p-values were calculated using Tukey HSD for all pairwise comparisons after one-way ANOVA (unequal variances) comparison. Linear and quadratic polynomial fits were performed in Origin 2020b.

We next evaluated if DAF-16 lifelong activity better quantifies other lifespan metrics. We compared DAF-16 total intensity to mortality quantiles (**Figure 3B**) at 50%, 75%, and 90%, which produced R^2^ of 0.48, 0.69, and 0.68 respectively. This suggests the DAF-16 response better captures when a *C. elegans* population perishes once it is already in decline or after 75% of the population has died. Previous studies indicate that DAF-16 has an important role in determining mortality and senescence in *C. elegans* (McElwee, Bubb and Thomas, 2003) and *D. Melanogaster* (Giannakou *et al*., 2007) at their late life stages. In light of this, our results would indicate that DAF-16 activity is relevant to determine mortality in old animals but not in the case of younger ones.

As mentioned previously, animals underwent DR repeatedly at days 1, 4, 7, and 10 of adulthood (**Figure 1B**). We observed that DAF-16 nuclear activity in animals previously exposed to DR returned to control levels after being well fed for 3 days. However, DAF-16 responsiveness to DR diminished at each following DR exposure (**Figure 3C**). It has been shown that animals that have been previously exposed to DR show increased resistance to stress (Smith *et al*., 2008). DAF-16 responsiveness could possibly be lower on repeated exposure to DR because the animals have already built resilience to stress from previous exposure. Alternatively, DAF-16 responsiveness could be reduced in aged worms, since the ability of animals to mount a response to stress has been shown to diminish with increasing age (Dues *et al*., 2016; Li *et al*., 2019). By day 10 of adulthood, once animals have passed their reproductive period, no significant change in DAF-16 activity was observed (**Figure 3C**).

We next analyzed the contributions of DAF-16 activity in specific days to mean lifespan (**Figure 3D**). Linear regressions of lifespan vs DAF-16 activity from individual days results in R^2^ of 0.71, 0.63, 0.75, and 0.45 at 1, 4, 7, and 10 days respectively. This analysis indicates that the DAF-16 response relevant for lifespan extension occurs during days 1, 4, and 7 for the DR regimes analyzed here. Likely, this results from a lack of responsiveness of DAF-16 at day 10. We analyzed the DAF-16 responsiveness from each day in all DR regimes and sorted the results by their corresponding mean lifespan (**Figure 3E**). In all cases, DAF-16 activity is significantly reduced at day 10, which occurs after animals stopped egg production (at days 7-8 of adulthood in our experiments). It has been shown that reproduction and lifespan are intertwined in *C. elegans* (Hsin and Kenyon, 1999; Luo and Murphy, 2011; Wang *et al*., 2014). Potentially, a loss of DAF-16 responsiveness could stem from animals turning-off prioritization of cell protection under stress after their reproductive period is over.

The regimes that result in the longest mean lifespan show the strongest DAF-16 response (6 and 12 hours in **Figure 3E**). Notably, the regime of 6 hours with 0 OP50 cells/ml exhibits the third longest mean lifespan but its DAF-16 response is low compared to the rest. Brief starvation has been shown to stimulate autophagy and prolong lifespan in *D. Melanogaster* (Eisenberg *et al*., 2014). Potentially, the lifespan extension observed in this particular regime is influenced to a lower extent by DAF-16 while other transcription factors, such as the autophagy modulator PHA-4 (Panowski *et al*., 2007; Hansen *et al*., 2008), play a larger role.

We next analyzed if total food availability could also be a good quantitative predictor of mean lifespan. We calculated the total amount of food available to animals up to day 11 of adulthood (end of the last exposure to DR), in other words we measured the areas under the curve for each DR regime in **Figure S4**. Fitting the data with a quadratic polynomial resulted in an R^2^ of 0.72 (**Figure 3F**), with the highest lifespan values corresponding to intermediate levels of food availability, as expected (Win *et al*., 2013). This analysis suggests that total food availability, rather than timing or level of food restriction, is more important in modulating lifespan. Presumably, DR regimes different than the ones explored in **Figure 1B**, but with similar areas under their curve could lead to equivalent mean lifespans.

### Intestinal cells and neurons have the largest DAF-16 activity and contribution lifespan

Since we observe DAF-16 in a variety of tissues, we then asked which cell-type contributes more to lifespan. Using machine learning, we developed a cell-type classifier to quantify endogenous DAF-16 activity with tissue-specificity. When adding the total intensity of nuclear DAF-16 of all cells of a given type, we observed that neurons and intestinal cells exhibited the largest responses (**Figure 4A**). Neurons reach a peak around 12 hours, while intestinal cells rapidly peak at 6 hours and remain mostly stable afterwards. Since we are quantifying total intensity from all nuclei of each cell type, both the number and size of the nuclei play a role in the comparison of DAF-16 tissue-specific activity. The higher response from neurons could stem from their higher number, with hundreds of neurons per animal as opposed to 20 intestinal cells. To account for this, we analyzed total intensity on a per cell basis. In this case, the largest individual contribution comes from intestinal cells (**Figure 4B**), albeit this result could be explained by intestinal cells being the largest cells in *C. elegans* (Zhang *et al*., 2017). This is confirmed by neurons having a higher mean intensity than intestinal cells (**Figure S5**). From this analysis, we can conclude that the lifespan extension exhibited by animals under the DR conditions analyzed comes mainly from DAF-16 activity in intestinal cells and neurons. This is further supported by the contributions of total intensity by cell type to mean lifespan (**Figure S6**), where we obtained linear regression R^2^ values of 0.78, 0.64, 0.47, and 0.00 for neuron, intestinal, muscle, and hypodermal cell types respectively. Note that this low R^2^ value for muscle cells is likely the result of an outlier corresponding with the control experiment. These results are in alignment with previous studies indicating that DAF-16 activity in the intestine of *C. elegans* increases the lifespans of *daf-16*(-) insulin/IGF-1-pathway mutants (Libina, Berman and Kenyon, 2003), and that insulin signaling in neurons is important to drive lifespan (Iser, Gami and Wolkow, 2007).

**Figure 4:**
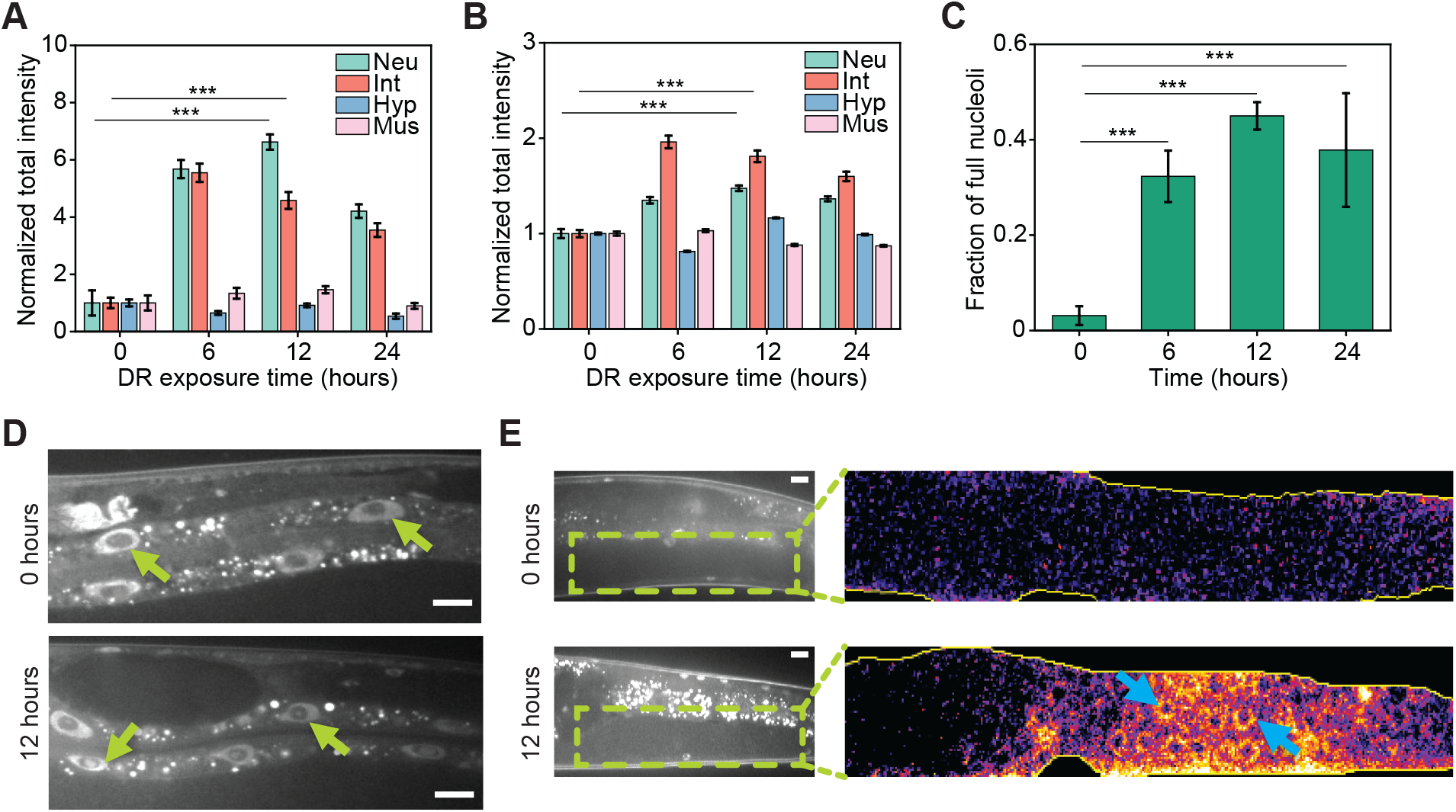
Tissue-specific analysis of DAF-16 reveals crucial role of intestine cells and neurons, nucleolar accumulation, and germline localization under liquid culture DR. A) DAF-16 normalized total intensity as a function of exposure time for various tissues. Neuron and intestine cells show the largest contribution to DAF-16 total intensity compared to hypodermis and muscle cells. B) DAF-16 total intensity per cell as a function of exposure time for various tissues. C) Fraction of intestinal nucleoli showing DAF-16 presence as a function of exposure times at a 10^8^ OP50 cells/ml food concentration. D) Examples of animals with empty nucleoli (0 hours) and DAF-16 in the nucleoli (12 hours) at a 10^8^ OP50 cells/ml food concentration. E) Inset highlights the gonad of a well-fed animal (0 hours) and one exposed to a food concentration of 10^8^ cells/ml (12 hours). DAF-16 in in the germline can be discerned in the DR-exposed animal (light-blue arrows). Error bars are SEM. p < 0.001 (***). All p-values were calculated using Tukey HSD for all pairwise comparisons after one-way ANOVA (unequal variances) comparison. Scale bars are 20 μM.

**DAF-16 shows activity in the germline and intestinal nucleoli**

During our experimental analysis, we observed that DAF-16 migration was not restricted to cell nuclei but also to the nucleolus in intestinal cells (**Figure 4C-D**). We observed that DAF-16 accumulates in 30-50% of intestinal nucleoli under the DR conditions used (**Figure 4C)**. This phenomenon was not observed with a transgenic strain that expresses DAF-16::GFP under the *daf-16* promoter, MAH97, developed with transgenic approaches that typically result in hundreds of copies of the reporter in a random genomic location. In these conditions, nucleolar localization is likely hidden due to the high brightness of such reporters, suggesting that DAF-16 activity in intestinal nucleoli can only be observed at endogenous levels of expression. The presence of nucleolar DAF-16 in the intestine could be an important player in the mechanisms of lifespan extension observed under DR, as it has been shown that the nucleolus plays an important role in lifespan extension (Kim *et al*., 2014; Tiku *et al*., 2016; Tiku and Antebi, 2018). Similarly, we observed DAF-16 in the germline of animals under DR (**Figure 4E**), although at extremely dim levels. This presence of DAF-16 was not observed in the MAH97 strain expressing DAF-16 tagged with GFP described above, potentially due to germline silencing of transgenic DNA (Kelly and Fire, 1998; Seydoux and Schedl, 2001; Aljohani *et al*., 2020). Tagging the endogenous DAF-16 locus could enable the reporter to bypass these silencing mechanisms.

## Discussion

In this work, we have utilized CRISPR/Cas9 and computer vision to quantitatively analyze the link between the endogenous spatiotemporal activity of the main IIS regulator, DAF-16, and longevity in *C. elegans*. We show that the transcription factor DAF-16 accounts by itself for 78% of the variability in lifespan observed in *C. elegans* under the DR regimes explored here. Previous studies have modulated lifespan by manipulating food concentration or intervening on the genetic pathways governing aging (Sagi and Kim, 2012; Hou *et al*., 2016). Our quantitative analysis reveals that modulation of a single transcription factor, through DR interventions, accounts for 78% of lifespan variability. Moreover, this work shows that this robust lifespan prediction is achieved by assessing endogenous DAF-16 spatiotemporal activity in a longitudinal manner and is directly affected by the food abundance throughout the animals’ life. Notably, the DAF-16 activity that is more relevant for lifespan extension occurred during *C. elegans* reproductive age, highlighting the close relationship between lifespan and reproduction observed in these animals (Hsin and Kenyon, 1999). We explored the activity of DAF-16 in different cell types, showing that intestinal cells and neurons have the largest contributions to DAF-16 activity leading to lifespan extension. This indicates that the mechanisms that govern DAF-16-dependent longevity in *C. elegans* under DR mainly take place in these cell types. This is emphasized by our finding of DAF-16 migrating into intestinal nucleoli, where some of these lifespan-modulating phenomena could be occurring.

Finally, our work opens the possibility of further exploration of other genetic pathways, transcription factors, and environmental factors in a holistic manner. For instance, it is unclear if the predictive power of endogenous DAF-16 activity holds under other environmental or genetic perturbations that also modulate lifespan. Moreover, determining whether the contributions of other longevity and stress signaling pathways are additive, or if these interact in a synergistic or antagonistic manner to determine lifespan will enable better understand longevity modulation in an integrative manner.

## Acknowledgements

*C. elegans* strains were provided by the CGC, which is funded by NIH Office of Research Infrastructure Programs (P40 OD010440). This work was supported in part by the U.S. National Institutes of Health (NIH) grants R00AG046911 and R21AG059099 and the NSF IOS grant 1838314.

## Author contributions

J.H. performed the experiments, developed image analysis algorithms, and analyzed data. J.H. and A.S.M. conceptualized the work and wrote the manuscript. A.S.M. supervised the study.

## Declaration of interests

The authors declare no competing interests.

## Notes

### Competing Interest Statement

The authors have declared no competing interest.

## References

Aljohani, M. D. et al. (2020) ‘Engineering rules that minimize germline silencing of transgenes in simple extrachromosomal arrays in C. elegans’, Nature Communications. Nature Publishing Group, 11(1), pp. 1–14. doi: 10.1038/s41467-020-19898-0.

Amrit, F. R. G. et al. (2014) ‘The C. elegans lifespan assay toolkit’, Methods, 68(3), pp. 465–475. doi: 10.1016/j.ymeth.2014.04.002.

An, J. H. and Blackwell, T. K. (2003) ‘SKN-1 links C. elegans mesendodermal specification to a conserved oxidative stress response’, Genes and Development. Cold Spring Harbor Laboratory Press, 17(15), pp. 1882–1893. doi: 10.1101/gad.1107803.

Bishop, N. A. and Guarente, L. (2007) ‘Two neurons mediate diet-restriction-induced longevity in C. elegans’, Nature. Nature Publishing Group, 447(7144), pp. 545–549. doi: 10.1038/nature05904.

Chen, J. and Caswell-Chen, E. P. (2003) ‘Why Caenorhabditis elegans adults sacrifice their bodies to progeny’, Nematology. Brill, 5(4), pp. 641–645. doi: 10.1163/156854103322683355.

Dickinson, D. J. et al. (2015) ‘Streamlined genome engineering with a self-excising drug selection cassette’, Genetics, 200(4), pp. 1035–1049. doi: 10.1534/genetics.115.178335.

Dues, D. J. et al. (2016) ‘Aging causes decreased resistance to multiple stresses and a failure to activate specific stress response pathways’, Aging, 8(4), pp. 777–795. doi: 10.18632/aging.100939.

Eisenberg, T. et al. (2014) ‘Nucleocytosolic depletion of the energy metabolite acetyl-coenzyme A stimulates autophagy and prolongs lifespan’, Cell Metabolism. Cell Press, 19(3), pp. 431–444. doi: 10.1016/j.cmet.2014.02.010.

Evans, T. (2006) ‘Transformation and microinjection’, WormBook. WormBook, pp. 1–15. doi: 10.1895/wormbook.1.108.1.

Ghosh, R. and Emmons, S. W. (2008) ‘Episodic swimming behavior in the nematode C. elegans’, Journal of Experimental Biology, 211(23), pp. 3703–3711. doi: 10.1242/jeb.023606.

Giannakou, M. E. et al. (2007) ‘Dynamics of the action of dFOXO on adult mortality in Drosophila’, Aging Cell. John Wiley & Sons, Ltd, 6(4), pp. 429–438. doi: 10.1111/j.1474-9726.2007.00290.x.

Greer, E. L. et al. (2007) ‘An AMPK-FOXO Pathway Mediates Longevity Induced by a Novel Method of Dietary Restriction in C. elegans’, Current Biology, 17(19), pp. 1646–1656. doi: 10.1016/j.cub.2007.08.047.

Greer, E. L. et al. (2011) ‘Transgenerational epigenetic inheritance of longevity in Caenorhabditis elegans’, Nature, 479(7373), pp. 365–371. doi: 10.1038/nature10572.

Greer, E. L. and Brunet, A. (2009) ‘Different dietary restriction regimens extend lifespan by both independent and overlapping genetic pathways in C. elegans’, Aging Cell. Blackwell Publishing Ltd, 8(2), pp. 113–127. doi: 10.1111/j.1474-9726.2009.00459.x.

Gruber, J. et al. (2009) ‘Deceptively simple but simply deceptive - Caenorhabditis elegans lifespan studies: Considerations for aging and antioxidant effects’, FEBS Letters, pp. 3377–3387. doi: 10.1016/j.febslet.2009.09.051.

Guthmueller, K. L., Yoder, M. L. and Holgado, A. M. (2011) ‘Determining genetic expression profiles in C. elegans using microarray and real-time PCR’, Journal of Visualized Experiments, 3791(53), pp. 1–7. doi: 10.3791/2777.

Han, S. K. et al. (2016) ‘OASIS 2: Online application for survival analysis 2 with features for the analysis of maximal lifespan and healthspan in aging research’, Oncotarget. Impact Journals LLC, 7(35), pp. 56147–56152. doi: 10.18632/oncotarget.11269.

Hansen, M. et al. (2008) ‘A role for autophagy in the extension of lifespan by dietary restriction in C. elegans’, PLoS Genetics, 4(2). doi: 10.1371/journal.pgen.0040024.

Henderson, S. T. and Johnson, T. E. (2001) ‘daf-16 integrates developmental and environmental inputs to mediate aging in the nematode Caenorhabditis elegans’, Current Biology, 11(24), pp. 1975–1980. doi: 10.1016/S0960-9822(01)00594-2.

Hou, L. et al. (2016) ‘A systems approach to reverse engineer lifespan extension by dietary restriction’, Cell Metabolism, 23(3), pp. 529–540. doi: 10.1016/j.cmet.2016.02.002.

Hsin, H. and Kenyon, C. (1999) ‘Signals from the reproductive system regulate the lifespan of C. elegans’, Nature. Nature Publishing Group, 399(6734), pp. 362–366. doi: 10.1038/20694.

Hsu, A. L., Murphy, C. T. and Kenyon, C. (2003) ‘Regulation of aging and age-related disease by DAF-16 and heat-shock factor’, Science, 300(5622), pp. 1142–1145. doi: 10.1126/science.1083701.

Iser, W. B., Gami, M. S. and Wolkow, C. A. (2007) ‘Insulin signaling in Caenorhabditis elegans regulates both endocrine-like and cell-autonomous outputs’, Developmental Biology. Academic Press, 303(2), pp. 434–447. doi: 10.1016/J.YDBIO.2006.04.467.

Kaeberlein, T. L. et al. (2006) ‘Lifespan extension in Caenorhabditis elegans by complete removal of food’, Aging Cell. John Wiley & Sons, Ltd, 5(6), pp. 487–494. doi: 10.1111/j.1474-9726.2006.00238.x.

Kelly, W. G. and Fire, A. (1998) ‘Chromatin silencing and the maintenance of a functional germline in Caenorhabditis elegans’, Development. The Company of Biologists, 125(13), pp. 2451–2456. doi: 10.1242/dev.125.13.2451.

Kenyon, C. J. (2010) ‘The genetics of ageing’, Nature, 464(7288), pp. 504–512. doi: 10.1038/nature08980.

Kim, Y. Il et al. (2014) ‘Nucleolar GTPase NOG-1 regulates development, fat storage, and longevity through Insulin/IGF signaling in C. elegans’, Molecules and Cells. The Korean Society for Molecular and Cellular Biology, 37(1), pp. 51–57. doi: 10.14348/molcells.2014.2251.

Kumsta, C. and Hansen, M. (2012) ‘C. elegans rrf-1 mutations maintain RNAi efficiency in the soma in addition to the germline’, PLoS ONE, 7(5). doi: 10.1371/journal.pone.0035428.

Kwon, E. S. et al. (2010) ‘A new DAF-16 isoform regulates longevity’, Nature, 466(7305), pp. 498–502. doi: 10.1038/nature09184.

Laranjeiro, R. et al. (2017) ‘Single swim sessions in C. elegans induce key features of mammalian exercise’, BMC Biology, 15(1), p. 30. doi: 10.1186/s12915-017-0368-4.

Lee, G. D. et al. (2006) ‘Dietary deprivation extends lifespan in Caenorhabditis elegans’, Aging Cell. John Wiley & Sons, Ltd, 5(6), pp. 515–524. doi: 10.1111/j.1474-9726.2006.00241.x.

Li, S.-T. et al. (2019) ‘DAF-16 stabilizes the aging transcriptome and is activated in mid-aged Caenorhabditis elegans to cope with internal stress’, Aging Cell. John Wiley & Sons, Ltd (10.1111), p. e12896. doi: 10.1111/acel.12896.

Libina, N., Berman, J. R. and Kenyon, C. (2003) Tissue-Specific Activities of C. elegans DAF-16 in the Regulation of Lifespan, Cell. doi: 10.1016/S0092-8674(03)00889-4.

Lin, K. et al. (2001) ‘Regulation of the Caenorhabditis elegans longevity protein DAF-16 by insulin/IGF-1 and germline signaling’, Nature Genetics, pp. 139–145. doi: 10.1038/88850.

Luo, S. and Murphy, C. T. (2011) ‘Caenorhabditis elegans reproductive aging: Regulation and underlying mechanisms’, Genesis. Wiley Subscription Services, Inc., A Wiley Company, pp. 53–65. doi: 10.1002/dvg.20694.

Masoro, E. J. (2005) ‘Overview of caloric restriction and ageing’, Mechanisms of Ageing and Development, pp. 913–922. doi: 10.1016/j.mad.2005.03.012.

Masoro, E. J., Yu, B. P. and Bertrand, H. A. (1982) ‘Action of food restriction in delaying the aging process’, Proceedings of the National Academy of Sciences of the United States of America, 79(13), pp. 4239–4241. doi: 10.1073/pnas.79.13.4239.

Mattison, J. A. et al. (2017) ‘Caloric restriction improves health and survival of rhesus monkeys’, Nature Communications, 8. doi: 10.1038/ncomms14063.

McElwee, J., Bubb, K. and Thomas, J. H. (2003) ‘Transcriptional outputs of the Caenorhabditis elegans forkhead protein DAF-16.’, Aging cell. John Wiley & Sons, Ltd, 2(2), pp. 111–121. doi: 10.1046/j.1474-9728.2003.00043.x.

Mitchell, D. H. et al. (1979) ‘Synchronous growth and aging of Caenorhabditis elegans in the presence of fluorodeoxyuridine’, Journals of Gerontology. Oxford Academic, 34(1), pp. 28–36. doi: 10.1093/geronj/34.1.28.

Panowski, S. H. et al. (2007) ‘PHA-4/Foxa mediates diet-restriction-induced longevity of C. elegans’, Nature, 447(7144), pp. 550–555. doi: 10.1038/nature05837.

Perez, A. R. et al. (2017) ‘GuideScan software for improved single and paired CRISPR guide RNA design’, Nature Biotechnology. Nature Publishing Group, 35(4), pp. 347–349. doi: 10.1038/nbt.3804.

Porta-de-la-Riva, M. et al. (2012) ‘Basic Caenorhabditis elegans methods: Synchronization and observation’, Journal of Visualized Experiments. MyJoVE Corporation, (64). doi: 10.3791/4019.

Portman, D. (2006) ‘Profiling C. elegans gene expression with DNA microarrays’, WormBook, pp. 1–11. doi: 10.1895/wormbook.1.104.1.

Possik, E. and Pause, A. (2015) ‘Measuring Oxidative Stress Resistance of Caenorhabditis elegans in 96-well Microtiter Plates’, Journal of Visualized Experiments, 99(99), pp. 527463791–52746. doi: 10.3791/52746.

Sagi, D. and Kim, S. K. (2012) ‘An engineering approach to extending lifespan in C. elegans’, PLoS Genetics, 8(6), p. 1002780. doi: 10.1371/journal.pgen.1002780.

Seydoux, G. and Schedl, T. (2001) ‘The germline in C. Elegans: Origins, proliferation, and silencing’, International Review of Cytology. Academic Press, 203, pp. 139–185. doi: 10.1016/S0074-7696(01)03006-6.

Sheaffer, K. L., Updike, D. L. and Mango, S. E. (2008) ‘The Target of Rapamycin Pathway Antagonizes pha-4/FoxA to Control Development and Aging’, Current Biology. Elsevier, 18(18), pp. 1355–1364. doi: 10.1016/j.cub.2008.07.097.

Smith, E. D. et al. (2008) ‘Age-and calorie-independent life span extension from dietary restriction by bacterial deprivation in Caenorhabditis elegans’, BMC Developmental Biology. BioMed Central, 8(1), pp. 1–13. doi: 10.1186/1471-213X-8-49.

Stiernagle, T. (2006) ‘Maintenance of C. elegans’, WormBook. doi: 10.1895/wormbook.1.101.1.

Stinchcomb, D. T. et al. (1985) ‘Extrachromosomal DNA transformation of Caenorhabditis elegans.’, Molecular and Cellular Biology. American Society for Microbiology, 5(12), pp. 3484–3496. doi: 10.1128/mcb.5.12.3484.

Stroustrup, N. et al. (2016) ‘The temporal scaling of Caenorhabditis elegans ageing’, Nature. Nature Publishing Group, 530(7588), pp. 103–107. doi: 10.1038/nature16550.

Tiku, V. et al. (2016) ‘Small nucleoli are a cellular hallmark of longevity’, Nature Communications. Nature Publishing Group, 8(1), pp. 1–9. doi: 10.1038/ncomms16083.

Tiku, V. and Antebi, A. (2018) ‘Nucleolar Function in Lifespan Regulation’, Trends in Cell Biology, pp. 662–672. doi: 10.1016/j.tcb.2018.03.007.

Wang, M. C. et al. (2014) ‘Gene Pathways That Delay Caenorhabditis elegans Reproductive Senescence’, PLoS Genetics, 10(12). doi: 10.1371/journal.pgen.1004752.

Wang, X. et al. (2009) ‘Identification of genes expressed in the hermaphrodite germ line of C. elegans using SAGE’, BMC Genomics, 10. doi: 10.1186/1471-2164-10-213.

Weindruch, R. (1996) ‘The retardation of aging by caloric restriction: Studies in rodents and primates’, Toxicologic Pathology. Taylor and Francis Ltd., 24(6), pp. 742–745. doi: 10.1177/019262339602400618.

Weinkove, D. et al. (2006) ‘Long-term starvation and ageing induce AGE-1/PI 3-kinase-dependent translocation of DAF-16/FOXO to the cytoplasm’, BMC Biology. BioMed Central, 4(1), pp. 1–13. doi: 10.1186/1741-7007-4-1.

Win, M. T. T. et al. (2013) ‘Validated liquid culture monitoring system for lifespan extension of Caenorhabditis elegans through genetic and dietary manipulations’, Aging and Disease. JKL International LLC, 4(4), pp. 178–185. Available at: /pmc/articles/PMC3733581/ (Accessed: 7 December 2021).

Zhang, N. et al. (2017) ‘The C. elegans intestine as a model for intercellular lumen morphogenesis and in vivo polarized membrane biogenesis at the single-cell level: Labeling by antibody staining, RNAi loss-of-function analysis and imaging’, Journal of Visualized Experiments. MyJoVE Corporation, 2017(128), p. 56100. doi: 10.3791/56100.

