## Supplementary Figures and Table for "Endogenous DAF-16 Spatiotemporal Activity Quantitatively Predicts Lifespan Extension Induced by Dietary Restriction"

### SUPPLEMENTARY FIGURES AND TABLES

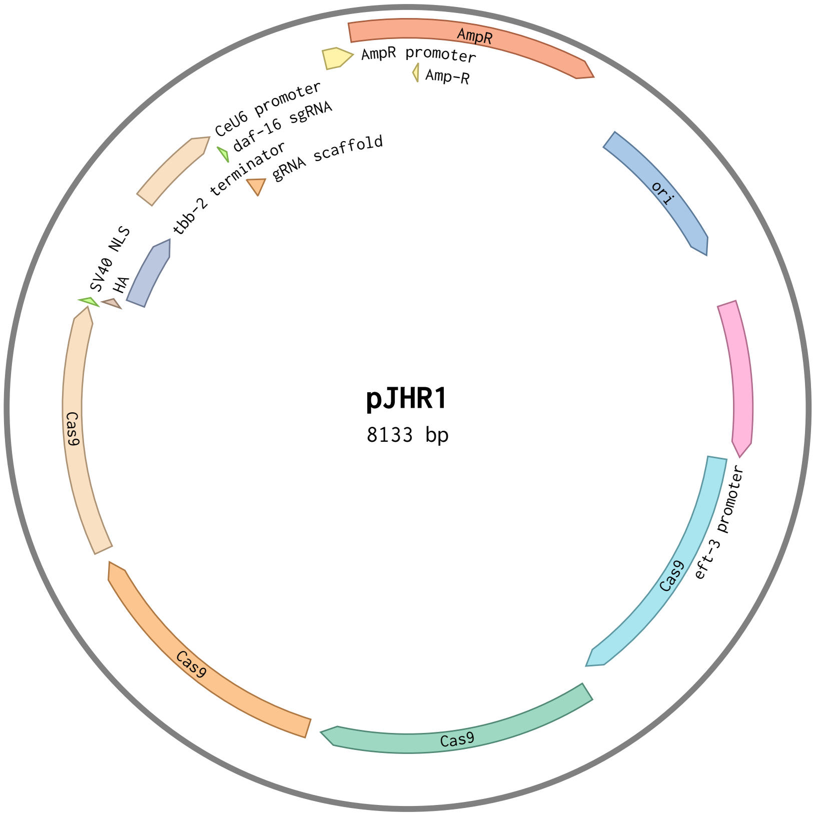


**Figure S1: Plasmid used for generation of transgenic strain with CRISP/Cas9.** pJHR1 contains the sgRNA targeting DAF-16 endogenous locus.

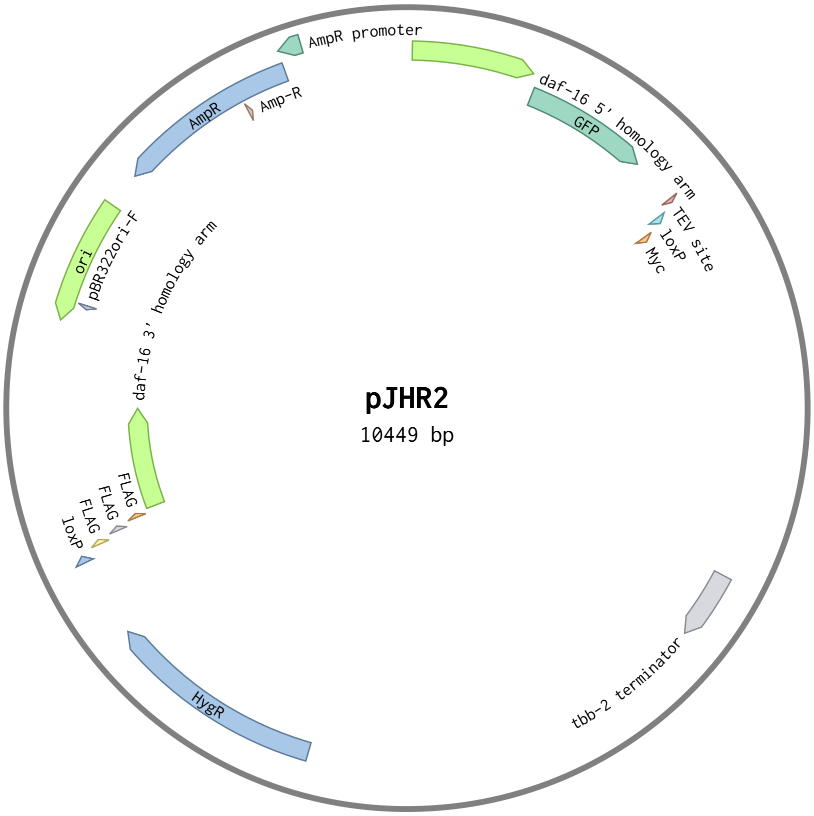


**Figure S2: Plasmid used for generation of transgenic strain with CRISP/Cas9.** pJHR2 contains GFP, self-excising cassette, and homology arms for DAF-16 tagging.

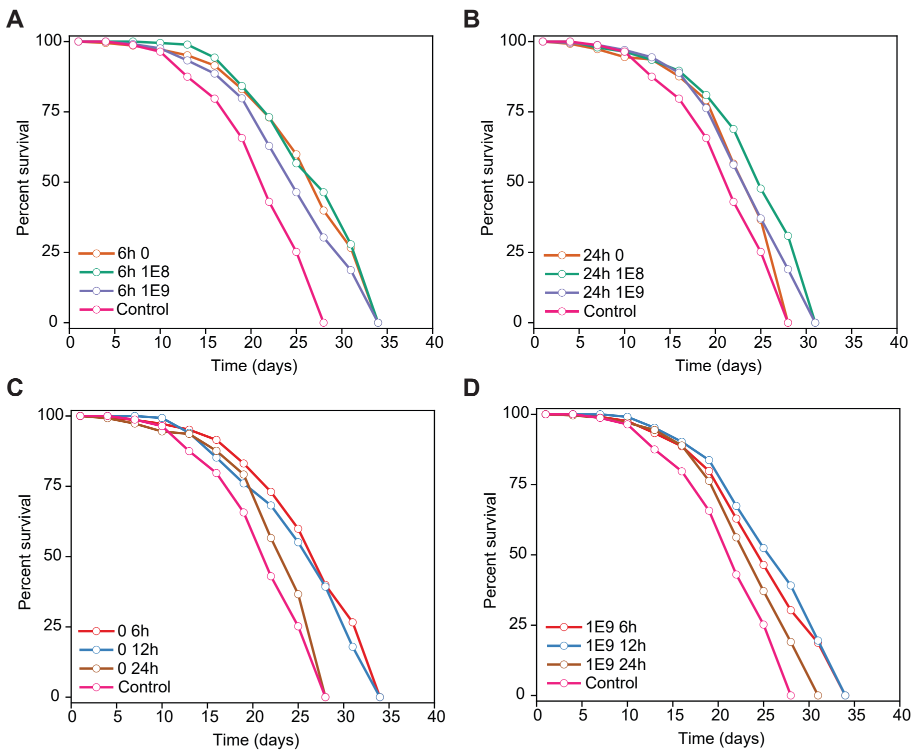


**Figure S3: Lifespan curves for various DR regimes.** A) Varying food concentration with 6 hours exposure time. B) Varying food concentration with 24 hours exposure time. C) Varying exposure time with a food concentration of 0 OP50 cells/ml. D) Varying exposure time with a food concentration of $10^9$ OP50 cells/ml.

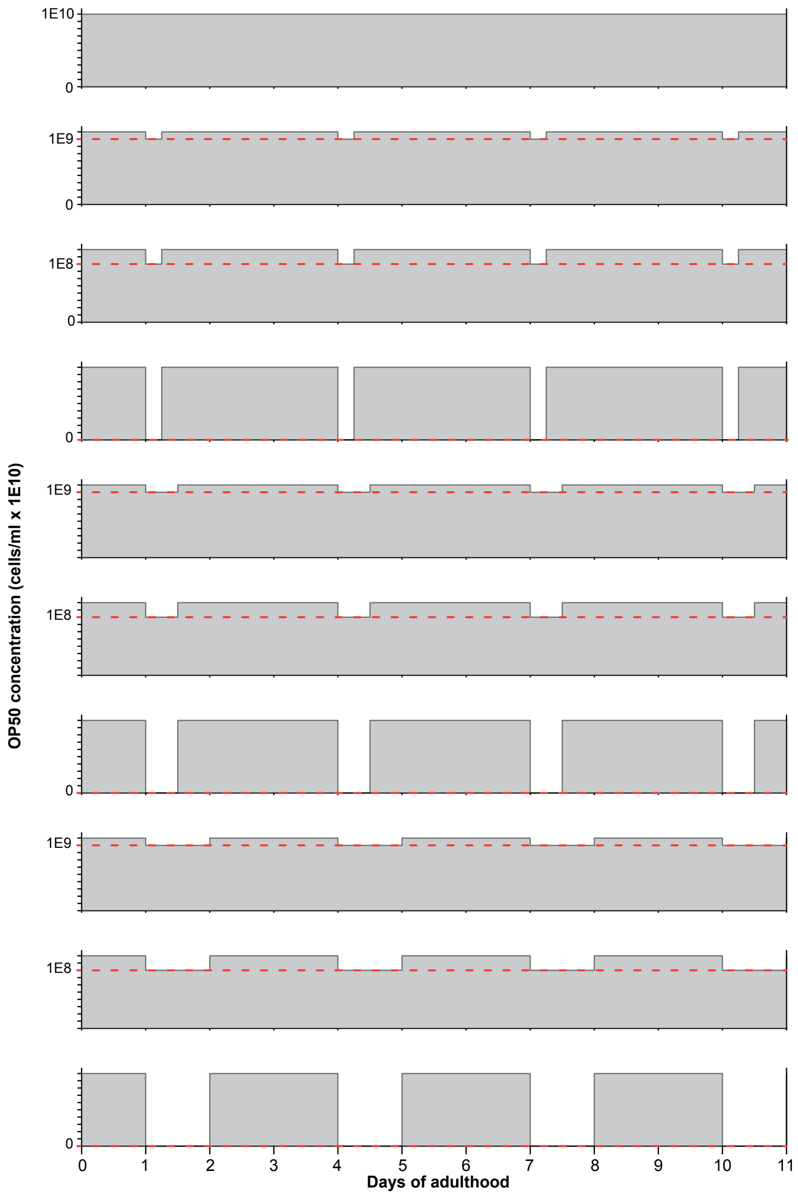


**Figure S4: Graphic representation of food availability for each of the DR regimes evaluated.** Grey areas were calculated for each DR regime and used in Figure 3F.

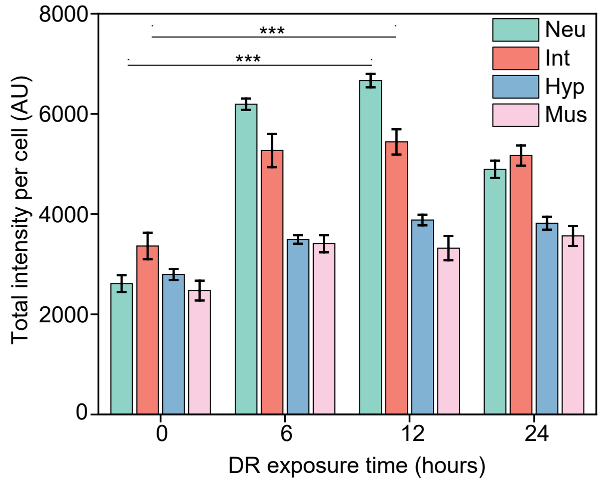


**Figure S5: Mean intensity per cell type at various exposure times.** Intestinal cells and neurons show the largest mean intensity compared to other cell types. Error bars are SEM. $p < 0.001$ (***). All p-values were calculated using Tukey HSD for all pairwise comparisons after one-way ANOVA (unequal variances) comparison.

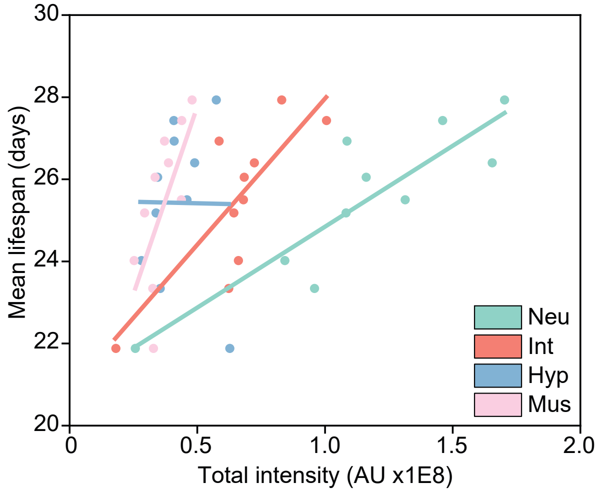


**Figure S6: Mean lifespan dependence on total intensity by cell type.** Intestinal cells and neurons show the largest contribution to mean lifespan. Linear fits were performed in Origin 2020b.

**Table 1:** Primers used for generation of transgenic line ASM10 *daf-16* (del2 [*daf-16*::GFP-C1^3xFlag]) I

| **Description** | **Primer** |
|---|---|
| 3' forward homology arm in pDD282 | CGTGATTACAAGGATGACGATGACAAGAGA TAAATTCTCTTCATTTTGTTTCCCC |
| 3' reverse homology arm in pDD282 | GGAAACAGCTATGACCATGTTATCGATTTCG GCTGTGATGATCGTTGAGTG |
| 5' forward homology arm in pDD282 | ACGTTGTAAAACGACGGCCAGTCGCCGGCA TACGGGCTCGATTTTCGTGA |
| 5' reverse homology arm in pDD282 | CATCGATGCTCCTGAGGCTCCCGATGCTCCC AAATCAAAATGAATATGCTGtCCT |
| sgRNA added to pDD162 | ATGAGCTGAGTCAAGCTGGA |
| Forward to introduce sgRNA in pDD162 | ATGAGCTGAGTCAAGCTGGAGTTTTAGAGC TAGAAATAGCAAGT |
| Reverse to introduce sgRNA in pDD162 | CAAGACATCTCGCAATAGG |
